# Appetite regulating genes in zebrafish gut; a gene expression study

**DOI:** 10.1101/2021.08.04.455091

**Authors:** Ehsan Pashay Ahi, Mathilde Brunel, Emmanouil Tsakoumis, Junyu Chen, Monika Schmitz

## Abstract

The underlying molecular pathophysiology of feeding disorders, particularly in peripheral organs, is still largely unknown. A range of molecular factors encoded by appetite-regulating genes are already described to control feeding behaviour in the brain. However, the important role of the gastrointestinal tract in the regulation of appetite and feeding in connection to the brain has gained more attention in the recent years. An example of such inter-organ molecular interaction can be the signals mediated by leptin, a key regulator of body weight, food intake and metabolism, with conserved anorexigenic effects in vertebrates. Leptin signal functions through its receptor (*lepr*) in multiple organs, including the brain and the gastrointestinal tract. So far, the regulatory connections between leptin signal and other appetite-regulating genes remain unclear, particularly in the gastrointestinal system. In this study, we used a zebrafish mutant with impaired function of leptin receptor to explore gut expression patterns of appetite-regulating genes, under different feeding conditions (normal feeding, 7-day fasting, 2 and 6-hours refeeding). We compared these expression patterns to those from wild-type zebrafish, in order to identify leptin-dependent differentially expressed genes located in the zebrafish gut. We provide evidence that most appetite-regulating genes are expressed in the zebrafish gut. On one hand, we did not observed significant differences in the expression of orexigenic genes after changes in the feeding condition, and only one orexigenic gene, *hcrt*, displayed differential expression under impaired leptin signal. On the other hand, we found 8 anorexigenic genes in wild-types (*cart2*, *cart3*, *dbi*, *oxt*, *nmu*, *nucb2a*, *pacap* and *pomc*), as well as 4 genes in *lepr* mutants (*cart3*, *kiss1, kiss1r* and *nucb2a*), to be differentially expressed in the zebrafish gut after changes in feeding conditions. Most of these genes also showed significant differences in their expression between wild-type and *lepr* mutant in at least one of the feeding conditions. Finally, we observed that impaired leptin signalling influences potential regulatory connections between anorexigenic genes in zebrafish gut, particularly connections involving *cart2*, *cart3*, *kiss1*, *kiss1r*, *mchr2*, *nmu*, *nucb2a* and *oxt*. Altogether, these transcriptional changes propose a potential role of the gastrointestinal tract in the regulation of feeding through changes in expression of certain anorexigenic genes in zebrafish.

## Introduction

In recent years, the prevalence of eating disorders has steadily increased in human populations worldwide [1]. The critical roles of various endocrine and neural factors are demonstrated in the regulation of feeding behaviour during eating disorders [2]. However, the underlying molecular pathophysiology of these disorders, particularly in peripheral organs, is still largely unknown. Feeding behaviour is regulated by specific regions in the brain, which control hunger-driven activities, through endocrine signals originated from the brain itself and peripheral organs [3, 4]. Factors secreted by the brain, in particular the hypothalamic area, and in other potentially important organs, control feeding behaviour by inhibiting (anorexigenic) or stimulating (orexigenic) appetite. Those key factors or appetite-regulating genes, which encode a range of neuropeptides and their cognate receptors, can be also classified in orexigenic and anorexigenic categories based on their functions [5, 6]. While most studies have focused on appetite-regulating factors in the brain, it is becoming increasingly evident that the gastrointestinal tract plays an important role in appetite and feeding regulation as well, and that bidirectional signalling between the gastrointestinal tract and the brain exists [7–11]. In addition, various metabolic changes causing eating disorders are influenced by factors produced by both organs [12], by gut microbiota [13, 14], as well as by homeostatic metabolite signals and nutrients [15, 16]. Altogether, these external and internal factors initially communicate with molecular signals within the gastrointestinal tissues in order to exert their effects along the gut-brain axis. Hence, signals mediated through the gastrointestinal tract are among the major contributors of peripheral mechanisms regulating appetite in the brain [17]. These signals can affect hypothalamic circuitry to determine cessation of energy intake during meal ingestion and the return of appetite and hunger after fasting [17].

A prominent example of these molecular signals linking multiple organs and conveying the nutritional status and the size of energy stores to the brain is the hormone leptin [18]. Leptin inhibits food intake and functions as an anorexigen during appetite regulation in several vertebrates, including fish [19], and loss of function mutations in orthologous leptin genes in zebrafish and mammals increases body weight and promotes obesity [20–22]. The conserved metabolic function of leptin signalling across vertebrates provides an opportunity to use zebrafish as model organism to explore molecular mechanisms underlying more specific leptin functions in appetite regulation and feeding. Unlike mammals, zebrafish contains two leptin hormone genes, *lepa* and *lepb*, expressed in several peripheral organs [23]. However, both leptin hormones mediate their signal through a single leptin receptor (*lepr*) gene, expressed in multiple brain regions, including the hypothalamus [24]. The regulatory effects of leptin signalling on appetite can be mediated through differential regulation of appetite regulating genes in the brain [25–28]. Nevertheless, leptin signal has also various effects on the gastrointestinal tract in relation to appetite and feeding [29, 30]. In mammals, leptin is produced in the stomach and its receptor is expressed in the intestine, mediating its effects on absorption of various nutrients in enterocytes, intestinal motility and cell proliferation, as well as secretion of different gastrointestinal hormones (reviewed in [30]). Subsequently, changes in intestinal conditions e.g. gut microbiota [14], trigger molecular signals, which are eventually sensed in specific brain regions. In zebrafish, both leptin and its receptor are expressed in the gastrointestinal tract and the enteric nervous system [31, 32]. The zebrafish intestinal expression of leptin and its receptor appeared to be responsive to fasting and feeding, indicating a potential regulatory role of leptin during changes of feeding conditions through the gut-brain axis [31, 32]. So far, studies addressing molecular mechanisms connecting the brain and gastrointestinal tissues under different feeding conditions are lacking in both fish and mammals. A first step to address this lack could be by generating information about transcriptional profiling of appetite regulating genes in the gut during changes in feeding conditions.

In this research, we assessed the expression levels of 37 appetite regulating genes in zebrafish gut (Table 1), which are already known to have also brain expression, under different feeding conditions [33]. The different feeding conditions to assess the expression dynamics of our target genes include normal feeding (control group), fasting and two refeeding steps (2 and 6 hours after refeeding). In addition, we investigated the expression of 11 genes with known gastrointestinal expression and function in zebrafish, as gut marker genes under the different conditions (Table 1). These marker genes were investigated to evaluate the effects of the different feeding conditions on gut functions. They include 4 genes for main anatomical compartments of gastrointestinal tract in zebrafish (*apoa1a*, *apoa4a*, *aqp3a* and *ctsl1*), 3 genes encoding important molecular transporters in gut (*pept1*, *slc2a5* and *sglt1*), 2 genes encoding regulators of the release of digestive enzymes (*ccka* and *cckar*), a target of leptin signal (*insa*) and a gut expressed orexigenic gene (*ghrl*) (see Table 1). To explore the role of leptin signalling, we compared the expression levels of all these genes between gut samples from wild-type zebrafish and zebrafish with impaired leptin receptor function (*lepr* mutant). To do this, we first validated a suitable reference gene with stable expression and used its expression as normalization factor against the expression of our target genes. Through this study, not only we profiled expression changes of appetite regulating genes in zebrafish gut, but also, found genes whose transcriptional changes could be under the influence of leptin signalling. Finally, through pairwise expression correlation analysis, we propose potential loss and gain of transcriptional regulatory links, between the appetite regulating genes under impaired leptin signal. Our study provides for the first time information about the expression dynamics of appetite-regulating genes in zebrafish gut, under both different feeding conditions and impaired leptin signalling.

**Table 1.**
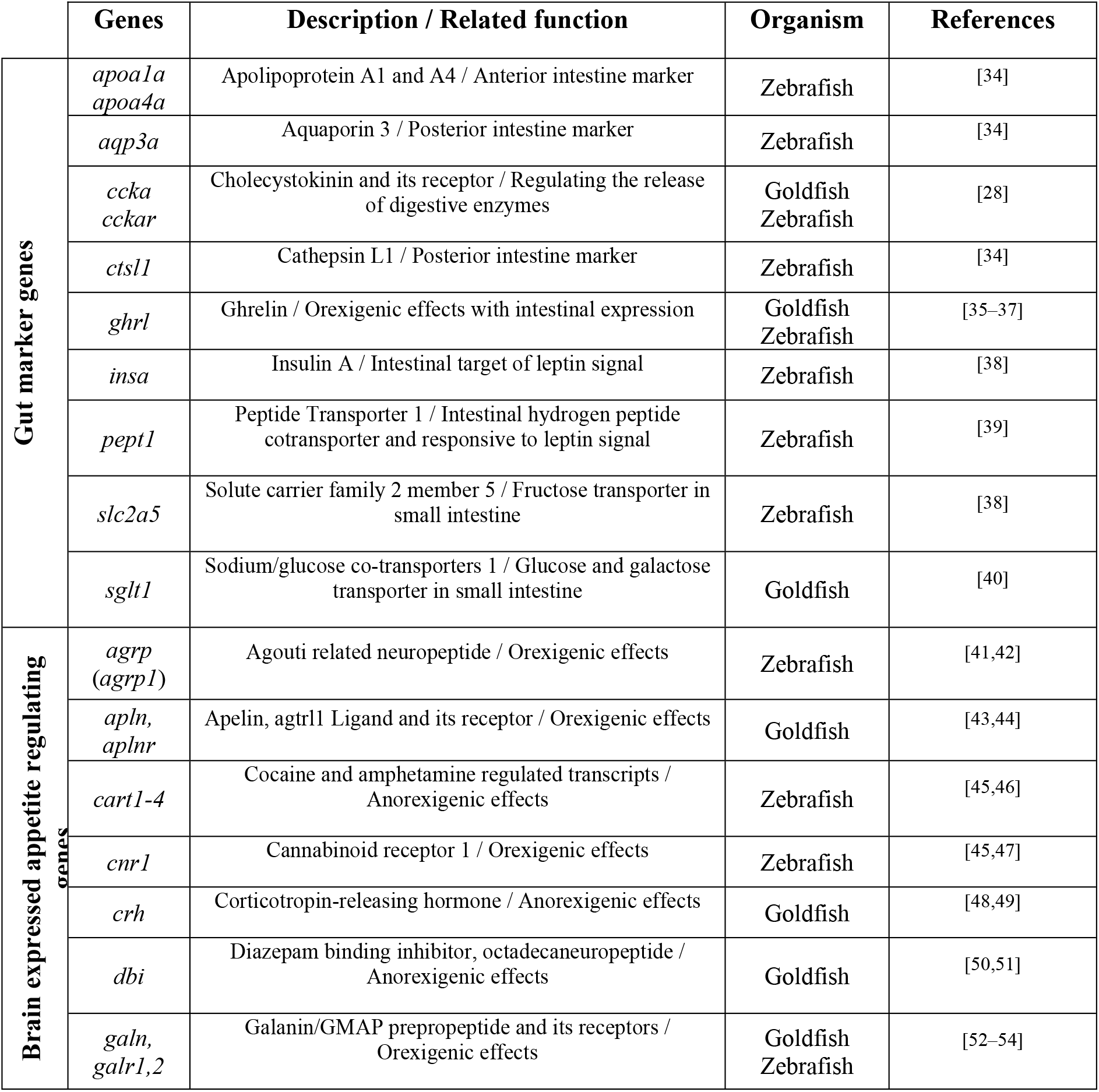
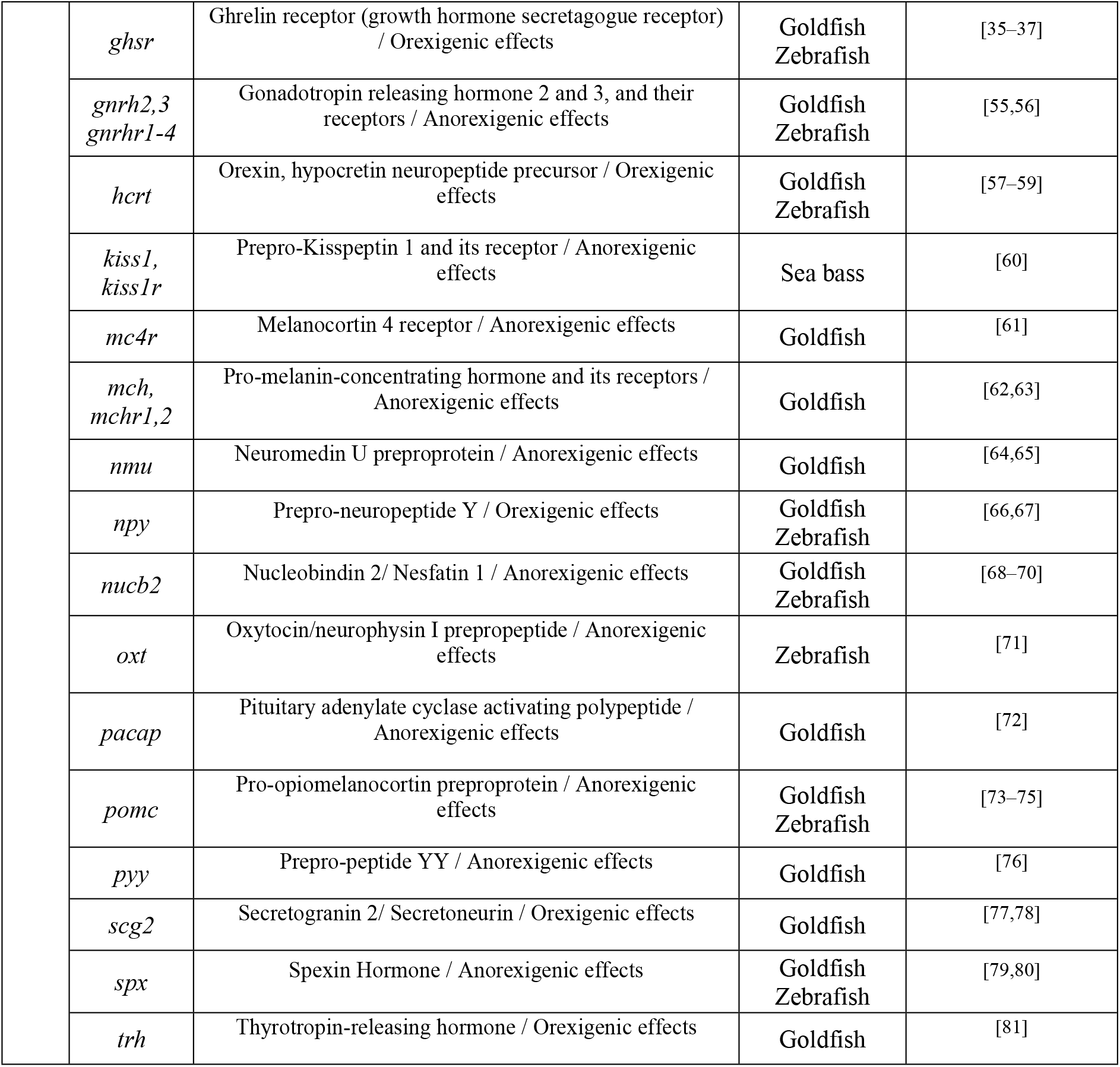
Candidate target genes studied and their related function in fish.

## Materials and Methods

### Fish breeding, fasting-refeeding scheme and sampling

*Lepr* sa12953 zebrafish were obtained from European Zebrafish Resource Centre. The mutation was created by the Sanger Institute for the Zebrafish Mutation Project, replacing a thymine with an adenine on chromosome 6 and causing the formation of a premature codon stop and an incomplete translation of *lepr* mRNA, resulting in a premature stop codon and a shortened polypeptide [82]. The fish were kept in 3-liter recirculating tanks at 28.4°C under light:dark conditions of 14:10 h at the SciLife lab Genome Engineering Zebrafish facility at Uppsala University. Water quality was checked regularly by the facility. The fish were fed with dry pellets once in the morning and with *Artemia* twice a day (middays and evenings). At the start of the experiment, 20 homozygote *lepr* mutant zebrafish and 20 wild-type zebrafish of similar age and size were divided into four experimental groups of 5 fish each, including both males and females. Fish in the control groups were fed according to the above-mentioned feeding schedule for 7 days; in the fasting groups, fish were fasted for 7 days and in the two refeeding groups, fish were fasted for 7 days and sampled 2 hours and 6 hours after refeeding. The control fish were sampled 2 hours after feeding. A detailed description of the experimental set up can be found in Ahi et al., (2019a). Fish were euthanized using a tricaine solution (MS-222) of 0.4 mg/ml and immersion in ice. The viscera of zebrafish were dissected and the entire gastrointestinal tract of each fish was carefully removed, washed in saline buffer and transferred into RNA*later*(Ambion Inc, Austin Texas) at 4°C for one day and then stored at -20°C. No significant differences in standard body length and net weight, as well as the hepato-somatic index (HSI) were shown between the genotypes (S1 Appendix). Fasting resulted in a 10%weight loss in both genotypes.

### RNA extraction and cDNA synthesis

We extracted the RNA of sampled guts, following the Trizol (Ambion) method. In brief, the dissected guts were removed from RNA*later*, dried and put into 1.5 ml Eppendorf tube with 200 μl of Trizol. Then, each sample was carefully homogenized with a 25G Terumo needle and BD Plastipak 1 mL syringe. The steps included addition of chloroform (Sigma-Aldrich), incubations, centrifugations, upper phase separation, addition of glycoblue (Ambion) and isopropyl alcohol (Sigma-Aldrich). RNA precipitation and ethanol (VWR) washes were followed, according to already described procedures [33]. The RNA pellets were dissolved in 10 μl of nuclease-free water (Ambion), and genomic DNA removal step was conducted using Turbo DNA-*free* kit (Ambion), according to the manufactureŕs instructions. The quality and quantity of RNA were assessed by NanoDrop 1000 v3.7 and 1000 ng of RNA input per sample was used for cDNA synthesis, with Superscript III First-Strand Synthesis System (Invitrogen). The cDNA synthesis protocol was performed in two steps, first by addition of 0.5 μl of random primers (50 ng/μl) and 0.5 dNTP (10 nM) to each RNA sample and incubation at 65°C for 5 minutes. Next, 2 μl of 5X First-Strand Buffer, 0.5 μl of DTT (0.1 M), 0.5 μl of RNase OUT (40 U/μl) and 0.5 μl of Superscript III RT (200 U/μl) were added to each RNA sample, followed by an incubation of 5 minutes at 25°C, 50 minutes at 50°C (RT enzyme reaction) and 15 minutes at 70°C (RT enzyme inhibition). The final volume of 10 μl of cDNA per sample was stored at -20°C until the qPCR step.

### Gene selection, primer design and qPCR

To find gene(s) with stable expression (reference gene(s)) across the experimental groups and genotypes, we chose 8 candidate genes, based on previous studies on zebrafish, in which suitable references genes were validated across different developmental stages, experimental conditions and tissues [83–85]. We then selected 37 candidate target genes involved in appetite regulation in studies on goldfish and zebrafish (Cypriniformes), as well as 11 marker genes with known functions in the gut, as summarized in Table 1. To design primers for qPCR, we first retrieved the gene sequences from a zebrafish database, http://www.zfin.org [86]. The sequences were then imported into CLC Genomic Workbench (CLC Bio, Denmark) and the exon/exon boundaries were specified, using the annotated *Danio rerio* genome available in the Ensembl database, http://www.ensembl.org [87]. The Primer Express 3.0 (Applied Biosystems, CA, USA) and OligoAnalyzer 3.1 (Integrated DNA Technology) software were used to design primers, as described in our previous study [33] (S1 Appendix).

The preparation of qPCR reactions was done by mixing 1 μl of diluted cDNA (1:20) from each sample with 7.5 μl of qPCR Master mix, PowerUp SYBR Green Master Mix (Thermo Fischer Scientific), 0.3 μl of forward and reverse primers (10 uM) and 6.2 μl of nuclease-free water. The MxPro-3000 PCR machine (Stratagene, La Jolla, CA) was used for quantification, combined with the MxPro software (Stratagene) for data analysis. We performed two technical replicates for each biological replicate (per gene), following the common sample maximization method [88], in order to optimize our assay in each run. The qPCR reaction program was set to 2 minutes at 50°C, 2 minutes at 95°C, and 40 cycles of 15 second at 95°C and 1 minute at 62°C. At the end of the amplification step, we performed a dissociation run (60°C – 95°C) and the threshold cycles and efficiencies were calculated. To calculate efficiency values (E) for each primer pair, we generated standard curves, by making serial dilution of pooled cDNA of pooled samples. The detailed analysis of standard curves and efficiencies are described in our previous study [33] (S1 Appendix).

### Gene expression analyses

In this study, we implemented three commonly used algorithms to validate the most suitable reference gene(s); BestKeeper [89], NormFinder [90] and geNorm [91]. These algorithms use different analysis methods to rank the most stably expressed reference genes, which their strengths and limitations are already discussed in details [92]. Next, the Cq value of the most stable reference gene, Cq _reference_, was used to normalize Cq values of target genes in each sample (ΔCq _target_ = Cq _target_ – Cq _reference_). Across all the samples, the biological replicate with the lowest expression for each gene was selected to calculate ΔΔCq values (ΔCq _target_ – ΔCq _calibrator_). The relative expression quantities (RQ values) was estimated by 2^−ΔΔCq^, and their fold changes (logarithmic values of RQs) were used for statistical analysis [93]. For the direct comparisons of gene expression levels between the genotypes (within each experimental condition), the student t-test was applied. On the other hand, to compare the expression dynamic of a target gene across the different conditions (within each genotype), analysis of variance (ANOVA) test was applied, followed by Tukey’s HSD post hoc tests. Benjamini-Hochberg method was also applied to correct for the false positive rate in the multiple comparisons [94]. Pairwise Pearson correlation coefficients between the appetite regulating genes were calculated within each genotype, in order to identify possible similarities in expression patterns. A dendrogram hierarchical clustering, using expression levels of the appetite regulating genes as input, was constructed, to represent similarities between the experimental conditions and the genotypes. R project (http://www.r-project.org) was used for the statistical analyses [95]. Finally, using a zebrafish database for protein interactome, STRING v10 (http://string-db.org/) [96], we investigated potential interactions between the gene products.

## Results

### Reference gene validation

Validation of stably expressed gene(s) in every specific experimental conditions (so-called reference genes) is an essential step, in order to calculate the relative gene expression levels of target genes [97–99]. In this study, we assessed the expression levels of 8 candidate reference genes in the gut of *lepr* mutant and wild-type zebrafish, under different feeding conditions. The expression levels of the candidate reference genes showed considerable variations, with highest expression (lowest Cq, Cq≈19) for *actb2*, and lowest expression (highest Cq, Cq≈30) for *tmem50a* (S1 Fig). Interestingly, the variations in the expression levels of the candidate reference genes in the gut were very similar to their expression variations in the brain [33]. All the three algorithms, BestKeeper, geNorm and Normfinder, ranked *tmem50a,* as the most stable reference gene (Table 2). In further analyses, we considered the expression level of *tmem50a* gene as a normalization factor to calculate the expression levels of our target genes in each sample.

**Table 2.** Ranking of candidate reference genes in zebrafish gut in wild-type and *lepr* mutant adults, under different feeding conditions. SD: standard deviation, r: Pearson product-moment correlation coefficient, SV: stability value, M: mean value of stability.

| BestKeeper |  |  |  | geNorm |  | NormFinder |  |
| --- | --- | --- | --- | --- | --- | --- | --- |
| Ranking | r values | Ranking | SD values | Ranking | M values | Ranking | S values |
| *tmem50a* | 0.928 | *ef1a* | 0.461 | *tmem50a* | 0.623 | *tmem50a* | 0.186 |
| *b2m* | 0.892 | *tmem50a* | 0.624 | *b2m* | 0.674 | *b2m* | 0.271 |
| *rpl13* | 0.882 | *actb2* | 0.626 | *ef1a* | 0.707 | *ef1a* | 0.276 |
| *ube2a* | 0.857 | *b2m* | 0.647 | *rpl13* | 0.732 | *rpl13* | 0.298 |
| *ef1a* | 0.823 | *rpl13* | 0.709 | *actb2* | 0.733 | *actb2* | 0.304 |
| *rplp0* | 0.819 | *p6gd* | 0.717 | *ube2a* | 0.783 | *p6gd* | 0.315 |
| *actb2* | 0.812 | *ube2a* | 0.799 | *p6gd* | 0.796 | *ube2a* | 0.359 |
| *p6gd* | 0.793 | *rplp0* | 0.944 | *rplp0* | 0.998 | *rplp0* | 0.401 |

### Expression analysis of gut marker genes in zebrafish

The first set of 11 target genes in this study are already known to be highly expressed in the gut and have different functions in the gastrointestinal system (Fig 1). We assessed the expression levels of these gut marker genes within wild-type and *lepr* mutant zebrafish, during the fasting-refeeding experiment. Among them, *apoa4a* and *slc2a5* genes showed expression differences between the feeding groups in both genotypes, whereas *aqp3a* only in wild-types, and *apoa1a* and *pept1* only in *lepr* mutants (Fig 1). The direct comparison between the two genotypes within each feeding group revealed expression differences only for *insa* in the control groups, *insa* and *apoa1a* in the fasting groups, and *sglt1* in the 6 hrs refeeding groups (Fig 2). Unexpectedly, we did not find any expression differences for *ghrl* in both genotypes and between different feeding conditions, which is an orexigenic gene with known expression in gut and brain [36]. The findings demonstrate that changes in feeding conditions can lead to expression differences in certain gut marker genes, although expression changes in only few of these genes are affected by impaired leptin signal in zebrafish gut.

**Figure 1.**
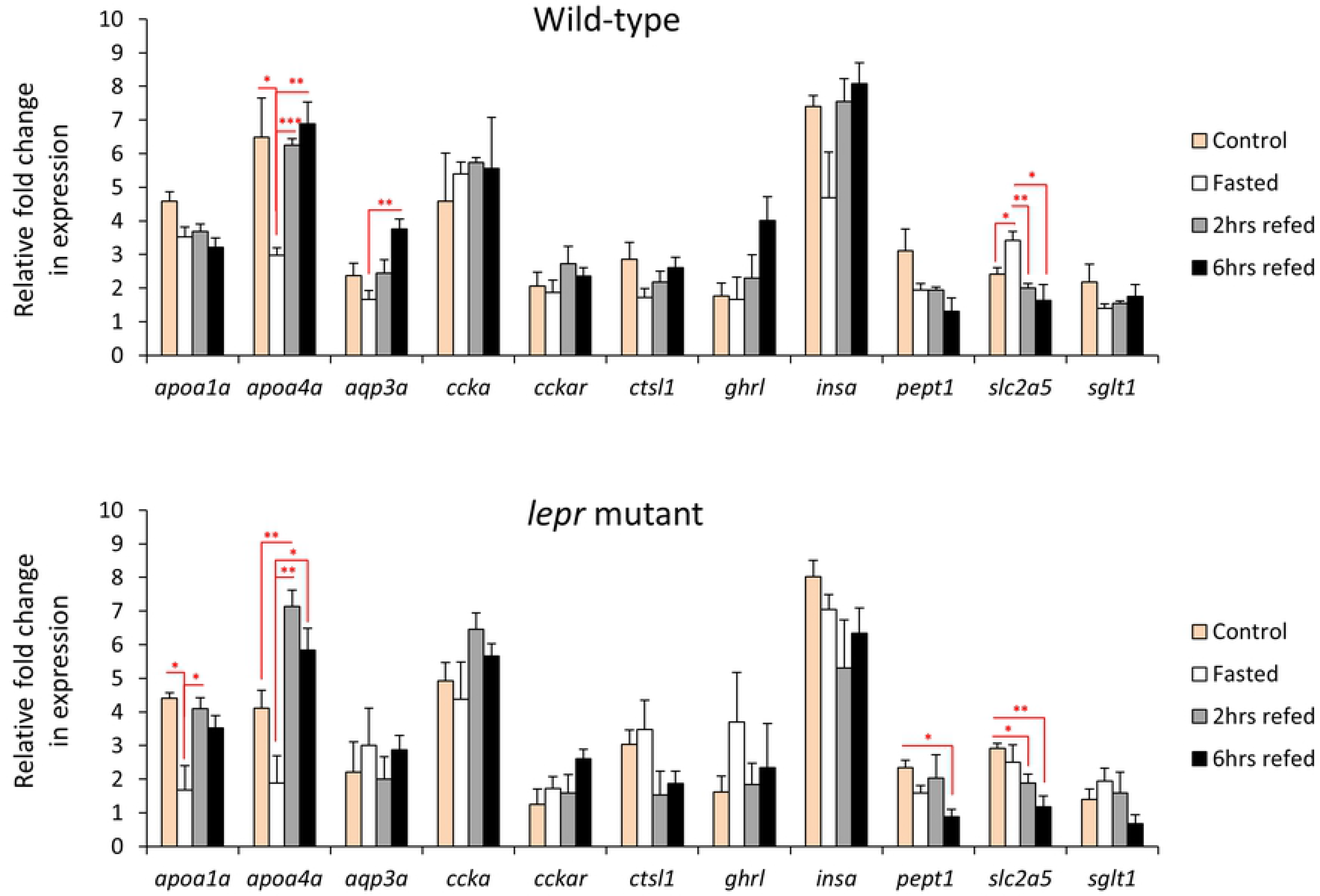
Expression dynamics of known gut-expressed genes in the gut of wild-type and *lepr* mutant zebrafish. Means and standard errors of fold changes in expression of five biological replicates are shown for each experimental group. Significant differences between the experimental groups in each genotype are delineated by asterisks (* *P* < 0.05; ** *P* < 0.01; *** *P* < 0.001).

**Figure 2.**
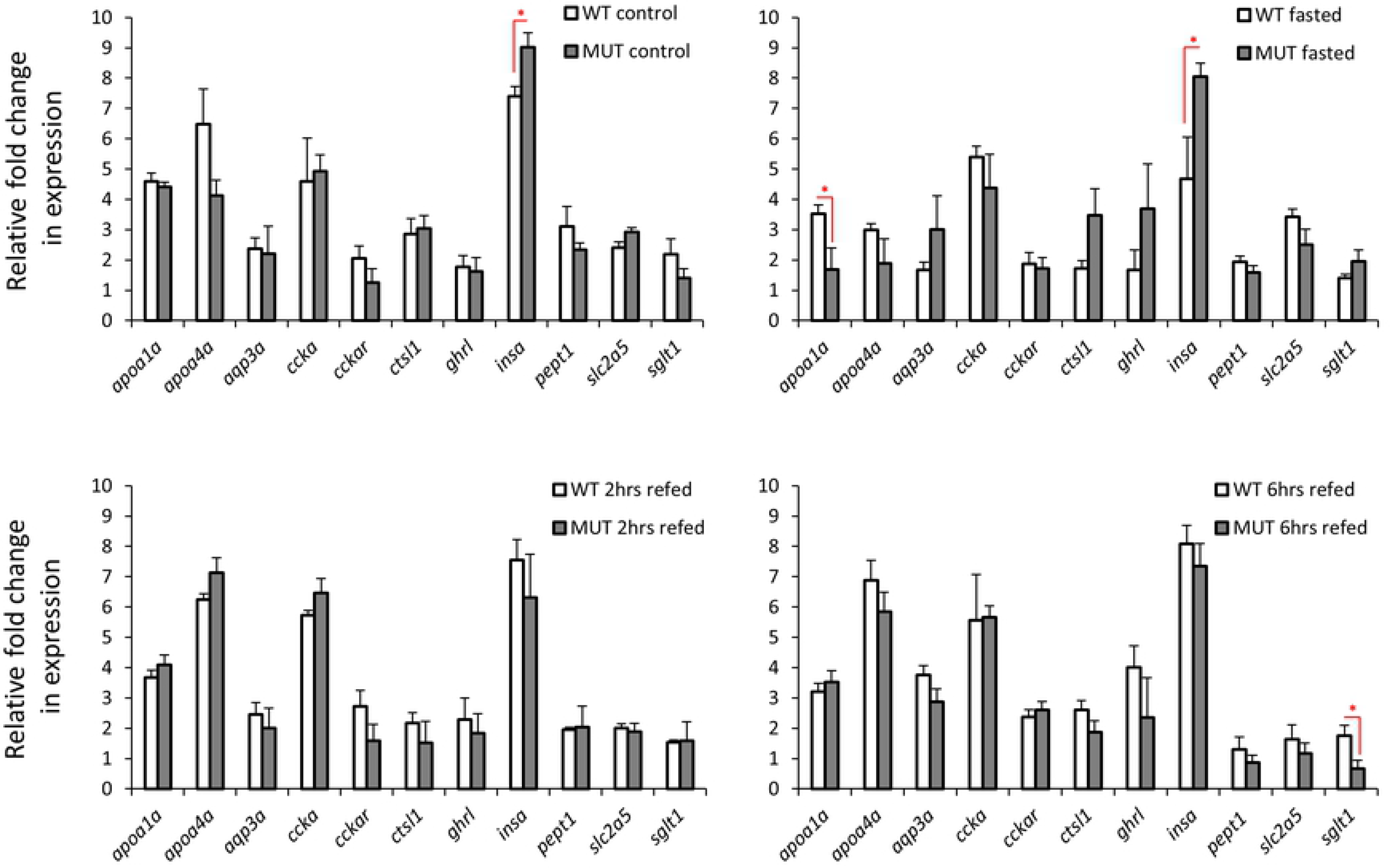
Expression differences of known marker genes in the gut of wild-type versus *lepr* mutant zebrafish in each feeding group. Means and standard errors of fold changes in expression of five biological replicates are shown for each experimental group. Significant differences between the *lepr* mutant and wild-type are delineated by asterisks (* *P* < 0.05).

### Expression analysis of orexigenic genes in zebrafish gut

We assessed the expression levels of 12 orexigenic genes, already known to have brain expression, within wild-type or *lepr* mutant gut during the fasting-refeeding experiment [33]. We found 8 of these genes to be expressed in the gut as well (Fig 3), however, unlike the brain, none of these genes showed expression differences between the feeding conditions in both genotypes. These observations indicate that the orexigenic genes expressed in the gut might not have functional roles during changes in feeding conditions, under both normal and impaired leptin signalling. Next, we compared the expression levels of the 8 orexigenic genes, between *lepr* mutant and wild-type for each feeding condition, and found only one gene, *hcrt*, showing increased expression in the mutants in the fasting groups (Fig 4). This might indicate a functional role of *hcrt* in zebrafish gut during fasting, which is controlled by leptin signal. In general, these results suggest minimal or absence of functional roles for the gut expressed orexigenic genes, under different feeding conditions and leptin signal in zebrafish.

**Figure 3.**
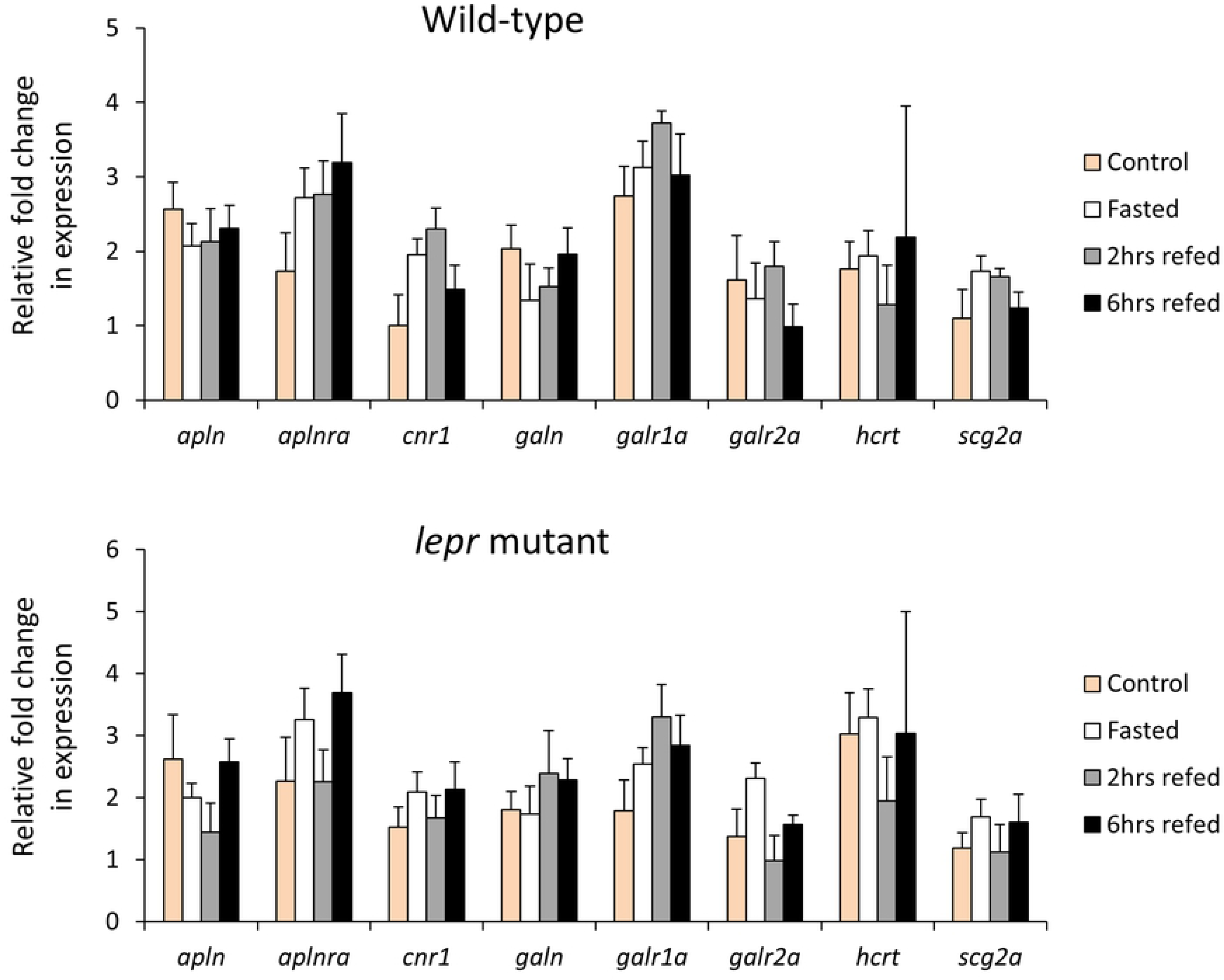
Expression dynamics of orexigenic genes in the gut of wild-type and *lepr* mutant zebrafish. Means and standard errors of fold changes in expression of five biological replicates are shown for each experimental group. No significant differences between the experimental groups were detected in both genotypes.

**Figure 4.**
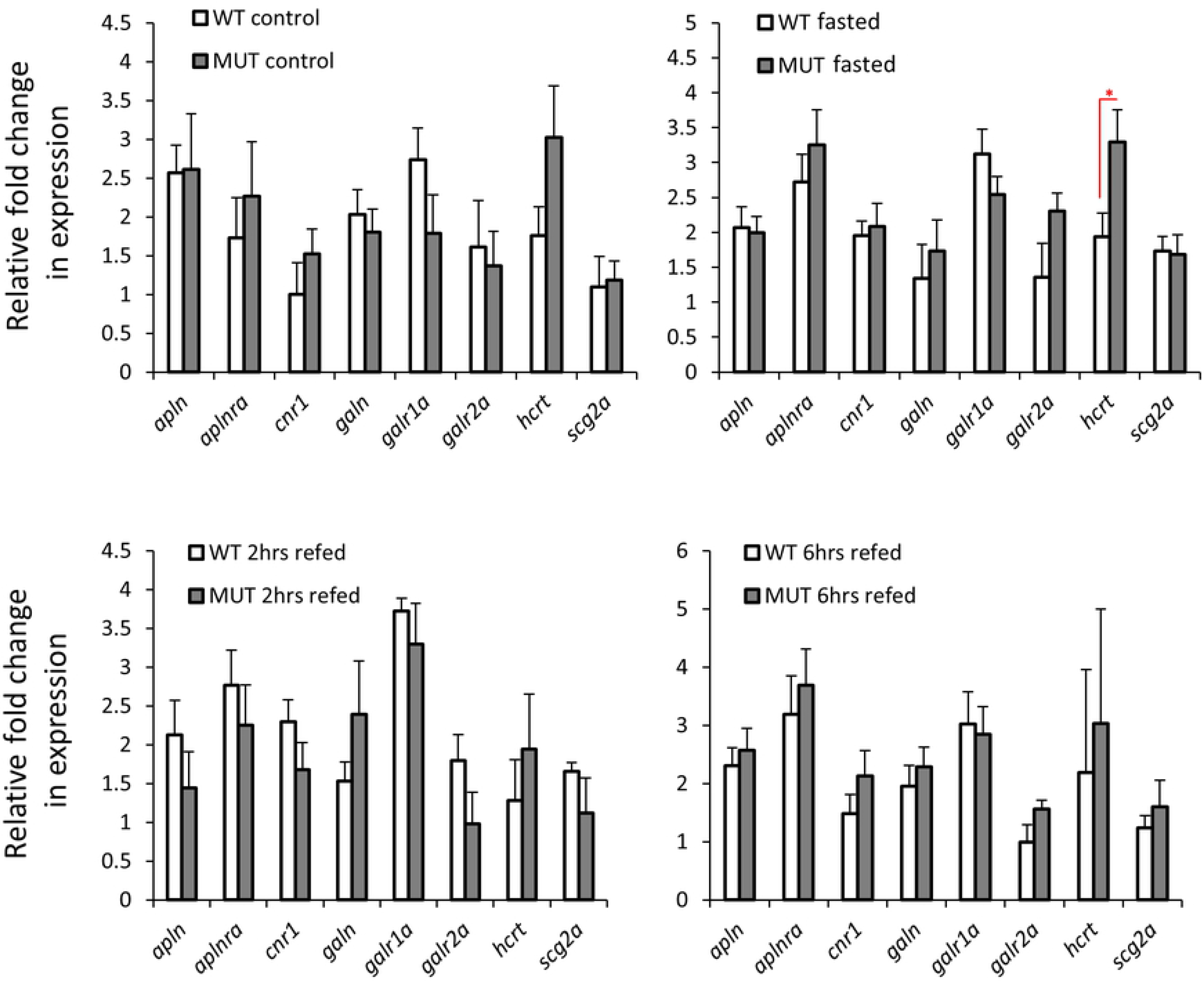
Expression differences of orexigenic genes in the gut of wild-type versus *lepr* mutant zebrafish in each feeding group. Means and standard errors of fold changes in expression of five biological replicates are shown for each experimental group. Significant differences between the *lepr* mutant and wild-type are delineated by asterisks (* *P* < 0.05).

### Expression analysis of anorexigenic genes in zebrafish gut

We assessed the expression levels of 25 anorexigenic genes, already known to have brain expression, in the gut of wild-type and *lepr* mutant zebrafish during the fasting-refeeding experiment [33]. We found 18 of these anorexigenic genes to be expressed in the gut as well (Fig 5). Among them, 8 genes showed expression differences between the feeding groups in the wild-type zebrafish (*cart2*, *cart3*, *dbi*, *oxt*, *nmu*, *nucb2a*, *pacap*, and *pomc*). This might indicate a functional role of the zebrafish gut in appetite control, through differential regulation of anorexigenic genes. The expression dynamics of these 8 genes showed also distinct variations, for example, *dbi* and *pomc* had increased expression during fasting, whereas *cart2*, *cart3* and *pacap* showed increased expression in at least one of the refeeding groups. In the *lepr* mutants, only 4 genes showed expression differences between the feeding groups, i.e. *cart3*, *kiss1* and *kiss1r* and *nucb2a* (Fig 5). This suggests that the expression differences of only 2 genes (*cart3* and *nucb2*) are independent of leptin signal, and that the disappearance of differences in the other genes can possibly be linked to functional leptin signal. Moreover, *kiss1* and *kiss1r* showed differences between the feeding groups only under the impaired leptin signal, indicating their potential leptin dependent function in the gut. In addition, we directly compared the expression of all the anorexigenic genes between the two genotypes, and observed expression differences for *mchr2* and *pomc* in control groups, *kiss1*, *kiss1r*, *nucb2a* and *pomc* in fasting groups, *dbi* in 2 hrs refeeding, and *kiss1*, *nmu* and *nucb2a* in 6 hrs refeeding groups (Fig 6). Genes *kiss1*, *nucb2a* and *pomc* showed higher expression in the wild-types in the fasting groups, but opposite expression differences (higher in the mutant) were evident in the control groups for *pomc* and in the 6 hrs refeeding groups for *kiss1* and *nucb2a*. Moreover, among the differentially expressed genes, *kiss1* and *kiss1r* had the strongest expression difference between the two genotypes, but in opposite manner (in fasting groups). Altogether, these findings suggest potential functional roles for certain gut expressed anorexigenic genes under different feeding conditions and leptin signal activities in zebrafish.

**Figure 5.**
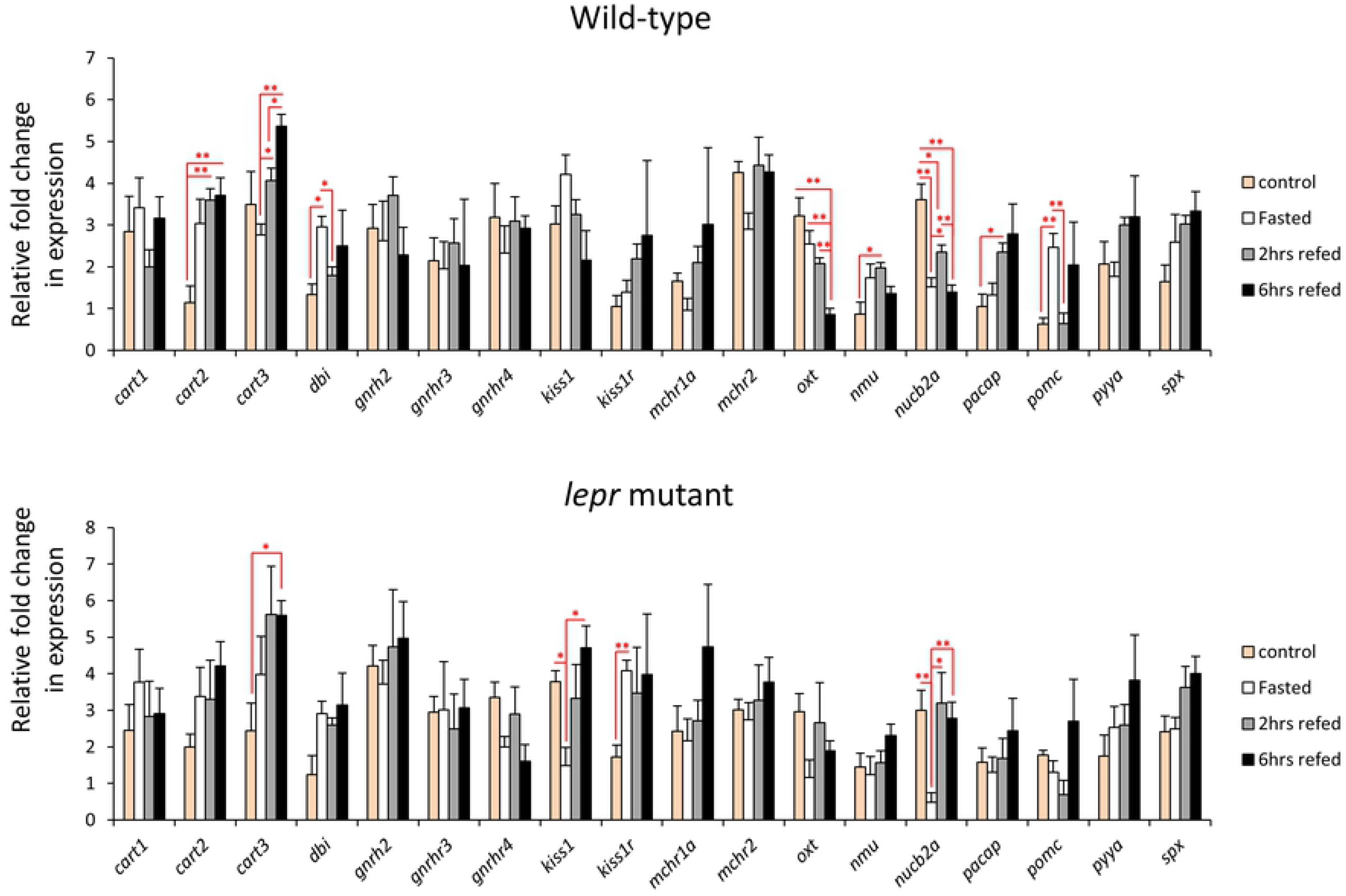
Expression dynamics of anorexigenic genes in the gut of wild-type and *lepr* mutant zebrafish. Means and standard errors of fold changes in expression of five biological replicates are shown for each experimental group. Significant differences between the experimental groups in each genotype are delineated by asterisks (* *P* < 0.05; ** *P* < 0.01).

**Figure 6.**
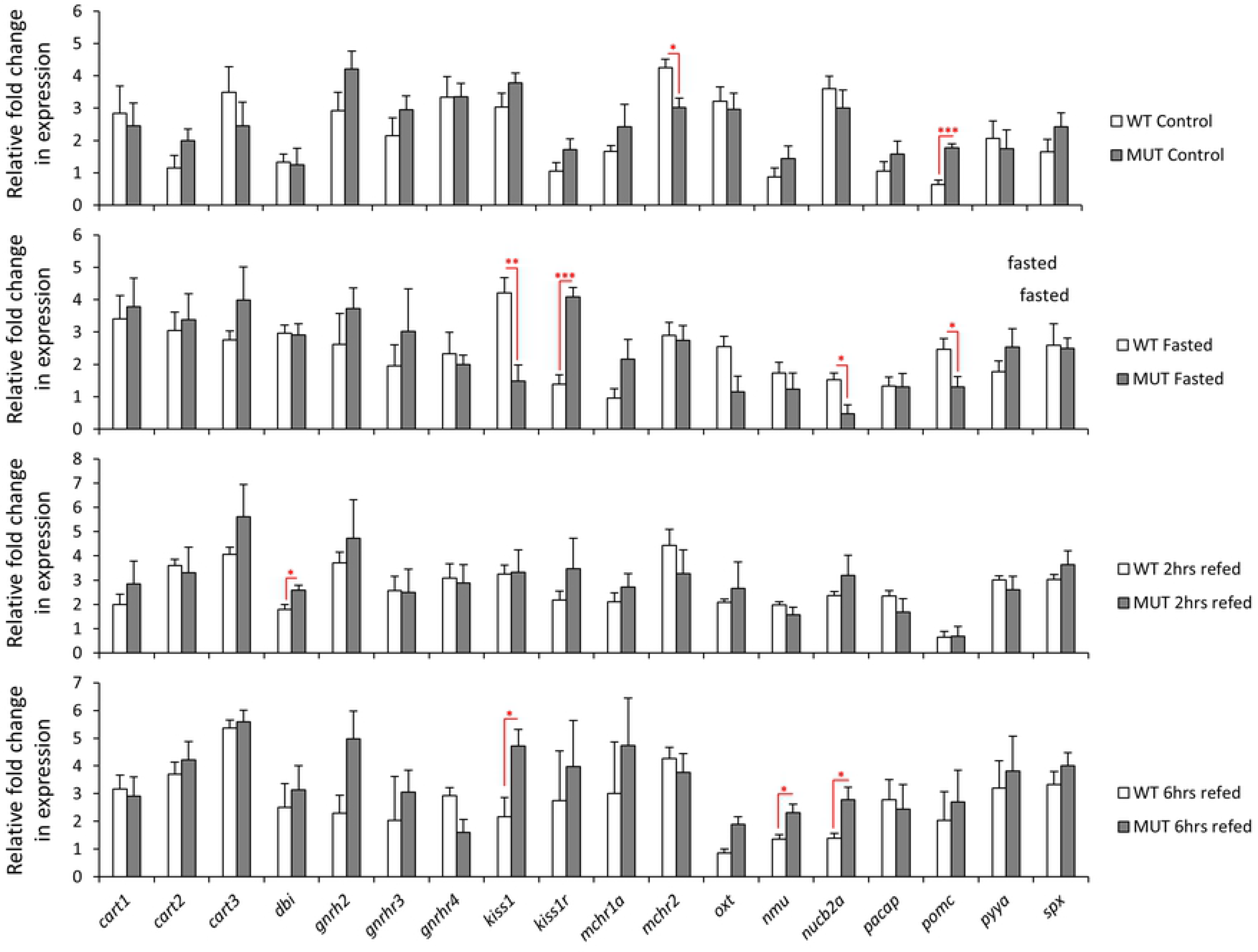
Expression differences of anorexigenic genes in the gut of wild-type versus *lepr* mutant zebrafish in each feeding group. Means and standard errors of fold changes in expression of five biological replicates are shown for each experimental group. Significant differences between the *lepr* mutant and wild-type are delineated by asterisks (* *P* < 0.05; ** *P* < 0.01; *** *P* < 0.001).

### Expression correlation analyses of the appetite regulating genes in zebrafish gut

In order to predict regulatory connections between the genes within each genotype, we performed pairwise analysis of expression correlation for the appetite-regulating genes, which showed differential expression in previous steps (Fig 7A). Most of the identified correlations were positive in both genotypes. Negative expression correlations were also found, but only in wild-type zebrafish and between *cart2*/*nucb2a*, *cart2*/*oxt*, *cart3*/*kiss1*, *mchr2*/*pomc* and *nmu*/*nucb2a*, indicating leptin-dependent negative transcriptional regulatory connections in zebrafish gut. Most of the positive expression correlations appeared to be present in both genotypes, suggesting their positive transcriptional regulatory connections independent of leptin signal in zebrafish gut. Although, few positive expression correlations appeared to be gained under impaired leptin signal in gut, i.e. between *cart2*/*kiss1r*, *cart3*/*mchr2*, *kiss1*/*mchr2* and *nmu*/*nucb2a*. Altogether, these results indicate transcriptional regulatory interactions between appetite-regulatory genes in the zebrafish gut. Moreover, these results revealed that impaired leptin signalling influences potential regulatory connections between appetite-regulating genes in the gut, particularly those connections, involving genes like *cart2*, *cart3*, *kiss1*, *kiss1r*, *mchr2*, *nmu*, *nucb2a* and *oxt*.

**Figure 7.**
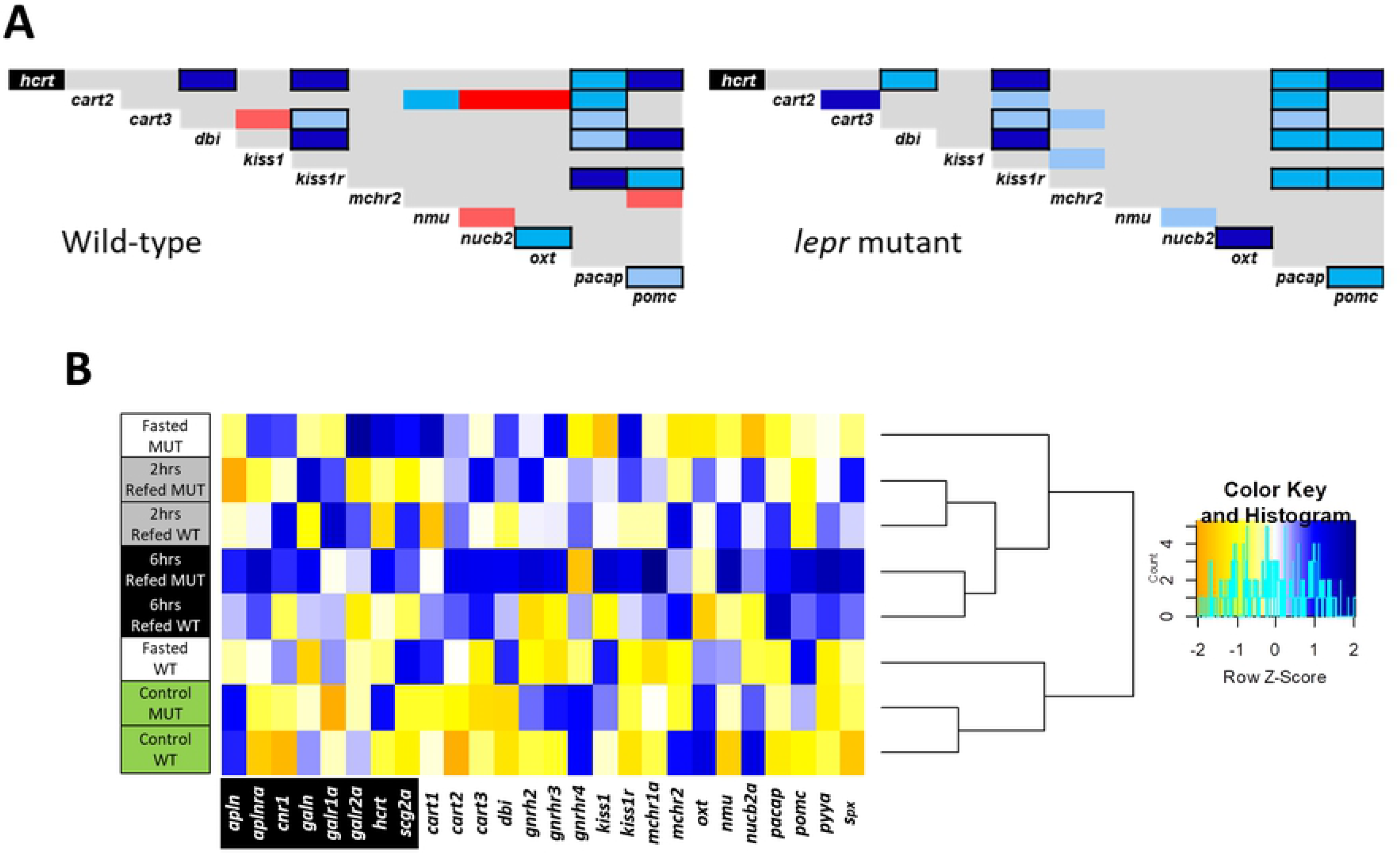
Connectedness of appetite-regulating genes and the feeding conditions based on the gene expression patterns. (**A**) Pairwise expression correlations of appetite regulating genes in the gut of wild-type and *lepr* mutant zebrafish. The blue and red colours respectively indicate positive and negative Pearson correlation coefficients and their light to dark shadings show significant levels of *P* < 0.05, *P* < 0.01 and *P* < 0.001, respectively. The gene specified with black background is an orexigenic gene whereas the rest are anorexigenic genes. Pairwise correlations delineated with black borders are similar between the two genotypes. (**B**) Clustering of the experimental conditions in each genotype based on similarities in expression patterns of the appetite regulating genes (the blue and yellow colors indicate higher and lower expression level, respectively).

In addition, using the same differentially expressed genes as input in a zebrafish protein interactome prediction tool (STRING v10, http://string-db.org/), we revealed potential molecular interactions between products of *hcrt*, *kiss1r*, *nmu*, *oxt*, *pacap* and *pomca*, as well as their transcriptional connections to *lepr* in zebrafish (S1 Fig). It also appeared that the product of *oxt* gene had highest number of molecular interactions with the rest of the gene products, as well as potential transcriptional connection to *lepr* in zebrafish.

Using the combined expression patterns of all appetite regulating genes expressed in zebrafish gut, we conducted hierarchical clustering of the experimental groups in both genotypes (Fig 7A). This led to identification of the similarities and divergences between the feeding conditions, which might also be affected by impaired leptin signal. The analysis revealed that the refeeding groups were clustered distantly from the control feeding groups in both genotypes, while the 2hrs and 6hrs refeeding groups were closely branched together for both genotypes (Fig 7B). On the other hand, the most distant branching between the two genotypes was observed for the fasting groups, i.e. the wild-type fasting group was more similar to the control group, whereas the mutant fasting group was clustered with the refeeding groups (Fig 7B). This indicates that the gut expression of appetite-regulating genes represents the strongest divergence between wild-types and *lepr* mutants during fasting.

## Discussion

The main aim of this study was to explore the expression dynamics of appetite-regulating genes in zebrafish gastrointestinal tract under different feeding conditions. We were also interested to investigate whether the impairment of leptin signal influences the expression of gut-expressed appetite regulating genes. Prior to addressing these aims, we first identified a reference gene, which is stably expressed in zebrafish gut under the different experimental conditions, and assessed the expression of a handful of gut marker genes (Fig 1 and 2). Two marker genes, *apoa4a* and *slc2a5*, showed changes in their expression dynamic in both wild-type and *lepr* mutant zebrafish. Apolipoprotein A4 gene (*apoa4a*) has conserved expression in villus-enterocyte of small intestine and hepatocytes of liver in zebrafish and mammals [34, 100]. Several functions have been attributed to *apoa4a*, such as regulating gluconeogenesis, lipid metabolism and transport, stomach emptying, acid secretion, and food intake (review [101]). In zebrafish, visceral upregulation of *apoa4a* is required for transition from endogenous to exogenous feeding, during mouth-opening larval stage [102]. The overexpression of *apoa4a* can lead to reduced food intake in both zebrafish and rats [103, 104]. In addition, it has been already demonstrated that extended fasting can reduce the expression of *apoa4a* in zebrafish liver [105] and its protein level in human body fluids [106]. Consistently, we found decreased expression of *apoa4a* in the gut in both genotypes, during fasting and this suggests its potential anorexigenic function in zebrafish. However, we found reduced expression of another apolipoprotein family member, *apoa1a*, only in the mutant group during fasting. Similarly, *apoa1a* is a conserved maker gene of the small intestine and liver in zebrafish and mammals [34, 107]. To our knowledge, there are no studies addressing transcriptional changes of *apoa1a* in response to fasting, its potential transcriptional regulation by leptin signal and its role in regulation of food intake in fish. Our findings of reduced *apoa1a* expression only in *lepr* mutant during fasting may imply on its leptin dependent role in lipid metabolism or food intake in zebrafish gut.

The second gene with differential expression dynamics during changes in feeding conditions in both genotypes, was *slc2a5*, but its expression pattern, unlike *apoa4a*, was different between the two genotypes. It displayed increased expression during fasting in the gut of wild-type zebrafish, whereas in *lepr* mutants, its expression was increased during normal feeding. Solute carrier family 2 member 5 or *slc2a5* (previously named *glut5*) encodes a fructose transporter responsible for fructose uptake by the small intestine [108, 109]. The reduction in intestinal expression of *slc2a5* during fasting, as well as its induced expression after refeeding, has been shown in rats [110]. Our finding show similar expression pattern for *slc2a5* in the zebrafish gut during fasting and refeeding. Interestingly, leptin signal has regulatory effects on hepatic expression of *slc2a5* in zebrafish [38]. In agreement with this study, we observed dysregulation of *slc2a5* in *lepr* mutants, indicating leptin-dependent transcriptional response of *slc2a5* to fasting and refeeding. The same study also showed increased visceral expression of an insulin gene (*insa* but not *insb*) in the absence of functional leptin signal [38]. Similarly, we also found higher expression of *insa* in the gut of *lepr* mutant zebrafish than wild-types in the control and fasting groups (Fig 2).

Two other marker genes showing differences in their expression dynamics between the genotypes were *aqp3a*, aquaglyceroporin-3, and *pept1*, intestinal H+/peptide cotransporter (also known as *slc15a1*) (Fig 1). *Aqp3a* showed increased expression levels at 6 hrs refeeding group compared to the fasting group, but only in wild-types. This indicates that this change might be dependent on leptin signal. *Aqp3a* encodes a conserved osmoregulatory channel protein on the membrane of epithelial cells in the large intestine in mammals and mucus cells of the posterior intestine of teleost fish [34, 111]. Thus, *aqp3a* is considered as a marker gene for the posterior part of intestine in vertebrates, which has an important role in water absorption, by increasing water permeability across cell membranes [34,111,112]. In mammals, leptin signal has been shown to control *aqp3* expression in liver and adipose tissue [113, 114]. The leptin dependent transcriptional regulation of *aqp3* is mediated through insulin and its downstream molecular cascade [113]. Such a regulatory connection between leptin signal and *aqp3a* has not been investigated in any tissues of teleost fish. Our finding suggests a regulatory link between leptin and *aqp3a* in zebrafish gut under different feeding conditions. The next gene, *pept1* encodes a major peptide transporter expressed in the proximal part of zebrafish intestine, is localized to the brush border membrane of the epithelial cells and mediates the uptake of di- and tripeptides from the lumen into the enterocytes [115]. In a previous study in zebrafish, *pept1* displayed increased expression during refeeding after short fasting [39], but we did not find such transcriptional changes. On the other hand, reduced intestinal expression of the mammalian orthologous of *pept1* has been reported in the absence of leptin signal [116]. In the present study, we found reduced *pept1* expression in 6 hrs refeeding group compared to the normal feeding group only in *lepr* mutant, which suggests potential regulatory connection between leptin and *pept1* expression in zebrafish gut. Finally, in the direct comparison between the two genotypes, we found reduced expression of *sglt1* in the mutants at 6 hrs refeeding group. The sodium/glucose co-transporter 1 gene or *sglt1* (also known as *slc5a1*) encodes an integral membrane protein, which is the major mediator of dietary glucose and galactose uptakes from the intestine [117]. A complex regulatory effect of leptin signal on intestinal expression of *sglt1* orthologue has been shown in mammals [118]. However, such a regulatory connection has not been investigated in fish.

None of the orexigenic genes showed any significant changes in their expression dynamics under the different feeding conditions in both genotypes. This could imply an absence of functional roles of orexigenic genes in feeding regulation in the zebrafish gut. An unexpected result was observed for *ghrl*, a gut expressed orexigenic gene, which is shown to be transcriptionally induced in response to fasting and reduced in response to refeeding in zebrafish [36], but we did not find any changes in both genotypes between different feeding conditions. In the direct comparisons between the two genotypes only one orexigenic gene, *hcrt* or *orexin*, showed higher expression in *lepr* mutants in the fasting groups and a similar tendency was also observed in the normal feeding groups (Fig 4). This suggests a repressive effect of leptin signal on *hcrt* expression in zebrafish gut during fasting. We have not detected such a regulatory connection in our previous study in zebrafish brain, indicating thus tissue specificity of the transcriptional regulation of *hcrt* by the leptin signal [33]. Hypocretin (orexin) neuropeptide precursor gene, *hcrt*, encodes a hypothalamic precursor protein that, after proteolytic processing, gives rise to two neuropeptides, orexin A and orexin B. These neuropeptides play role in various physiological processes, such as regulation of sleep-wake cycle, reward-seeking, addiction, stress and feeding behaviour in zebrafish [119]. During fasting, *hcrt*, acts as a potent orexigenic neuropeptide in zebrafish brain [59]. In goldfish, leptin signal represses *hcrt* brain expression and thus inhibits its orexigenic effect on feeding behaviour [25].

Unlike the orexigenic genes, almost half of the gut-expressed anorexigenic genes (8 out of 18 genes) showed expression differences during changes in feeding conditions (Fig 5). This suggests that the potential regulatory effects of the gut on feeding regulation are mediated through anorexigenic genes in zebrafish. Among these differentially expressed anorexigenic genes, only two, *cart3* and *nucb2a*, displayed changes in their expression dynamics in the gut of *lepr* mutants. This indicates that leptin signal plays important regulatory roles in transcriptional changes of the other 6 anorexigenic genes (*cart2*, *dbi*, *oxt*, *nmu*, *pacap* and *pomc*) in the zebrafish gut. For instance, induced expression of *cart2*, a member of cocaine and amphetamine regulated transcripts genes, in the refeeding groups, was only observed in the gut of wild-type zebrafish. This finding is opposite to their expression in the brain, where all members of *cart* genes showed reduced expression during fasting and refeeding in the wild-type zebrafish [33]. This also suggests that a recently found gene regulatory network (GRN) in zebrafish brain, consisting leptin signal and *sp1* transcription factor as upstream regulators of cart genes, i.e. *lepr-sp1-cart* axis, does not exist in the gut [120]. *Cart* genes are shown to have conserved anorexigenic role in teleosts [28], including zebrafish [45, 46]. Although regulatory connections between *cart* genes and leptin are found in other species, such as goldfish [121], and mammals [122, 123], these connections are only reported in the brain.

Perhaps the most striking finding among the anorexigenic genes was the differential expression of prepro-kisspeptin 1, *kiss1,* and its receptor, *kiss1r*, between the different feeding conditions, only in the *lepr* mutant fish, as well as between the genotypes during fasting (Fig 5 and 6). Although, *kiss1* and *kiss1r* are expressed in zebrafish gut and brain [124], so far their anorexigenic roles have only studied in sea bass (a Perciforme species) [60]. In mammals, the brain expression of *kiss1* is induced by leptin administration [125], while fasting reduces its brain expression levels [126, 127]. Consistently, we have already observed reduced *kiss1* expression after fasting in the brain of wild-type zebrafish but not in the *lepr* mutant, indicating its leptin-dependent transcriptional regulation during fasting in zebrafish [33]. In the gut, however, we found an opposite pattern; where *kiss1* expression was reduced during fasting only in the *lepr* mutants and not in the wild-type fish. This suggests a potential opposite regulatory function of leptin signal on *kiss1* expression between brain and gut in zebrafish. Interestingly, the kisspeptin receptor, *kiss1r*, which showed no expression difference between the genotypes in the brain [33], displayed an opposite expression pattern to its ligand gene (*kiss1*) in the gut, with increased expression during fasting. The regulatory connection between *kiss1* (and its receptor) and leptin signal is not studied in the digestive system of vertebrates and these observations stress the necessity of future functional studies in the gut to unravel potential links. These results also suggest that the functional role of leptin signalling through regulation of *kiss1/kiss1r* during fasting/refeeding might be more important in the gut than in the brain of zebrafish.

Another interesting finding was the increased expression of *pomc*, pro-opiomelanocortin gene, during fasting in the gut of wild-type zebrafish and the absence of these changes in the *lepr* mutant fish. The basal gut expression level of *pomc* was higher in the *lepr* mutant under normal feeding condition, which was opposite to our previous findings in the brain [33]. During fasting, however, this pattern was reversed in the gut, with higher expression in wild-type, while in our previous study, no difference was observed between the two genotypes in the brain during fasting [33]. The implications of these changes are difficult to interpret, but it is obvious that there is a distinct leptin-dependent transcriptional regulation of *pomc* between the two organs under similar feeding conditions.

In zebrafish brain, almost all of the anorexigenic genes regulated by leptin signal showed reduced expression in the *lepr* mutant under normal feeding condition [33]. In the direct comparisons between the two genotypes in the gut, *mchr2* was the only anorexigenic gene with reduced expression in the *lepr* mutants under normal feeding condition (Fig 6). *Mchr2* encodes a melanin concentrating hormone (MCH) receptor gene and although the role of MCH signal in appetite-regulation has not been investigated in zebrafish, its anorexigenic role is already demonstrated in goldfish [62, 63]. In mice, leptin acts as an upstream regulator of MCH signal in the brain [128, 129]. Apart from the role of MCH signal in the regulation of immune responses, no studies have addressed other potential functions of this signal in zebrafish gut. Nucleobindin 2a or nesfatin 1, *nucb2a*, was another gene showing differences in the direct comparisons between the two genotypes, however, these differences were in opposite direction, i.e. reduced expression during fasting, but increased expression in 6 hrs refeeding in the gut of *lepr* mutant zebrafish. The anorexignenic role of *nucb2a* is already described in zebrafish, and interestingly, fasting results in decreased expression, both in the brain and the gut [70]. This may indicate that the decreased expression of *nucb2a* in both organs mitigates its anorexigenic effects during fasting in zebrafish. Our findings confirm the reduced expression of *nucb2a* during fasting in both genotypes, indicating a leptin independent regulation. However, the increase in *nucb2a* expression during refeeding, only in *lepr* mutants, might be linked to the function of leptin. Finally, *dbi* and *nmu* showed higher expression in the *lepr* mutants than wild-types in one of the refeeding groups. The octadecaneuropeptide or ODN is an endozepine peptide, which is generated by the cleavage of the product of *dbi* gene (diazepam-binding inhibitor) in the brain. In goldfish, brain injection of ODN has been shown to inhibits food intake and locomotor activity [50, 51]. These anorexigenic effects in goldfish are mediated through a *mc4r* and *crh* dependent signals [51], but we have not detected the expression of neither *mc4r* nor *crh* in the zebrafish gut. This suggests that the potential function of ODN is mediated through different signals in zebrafish gut.

Neuromedin U preproprotein gene, *nmu*, encodes a multifunctional neuropeptide, which controls the circulatory and digestive systems, energy homeostasis, and acts as anorexigenic factor in goldfish [64, 65]. mRNA transcripts of *nmu* were found in both the brain and the gut of goldfish [64], and in the brain of zebrafish [33]. Moreover, brain injection of Nmu causes suppression of food intake and locomotor activity in a dose-dependent manner in goldfish [64]. Although, *nmu* encoded protein is considered as a brain-gut neuropeptide, its exact role in feeding regulation through the digestive system remains unclear. The anorexigenic effects of *nmu* are mediated through *crh* or corticotropin-releasing hormone receptor in the goldfish brain [65], and we also found *crh* to be a downstream target of leptin signal in zebrafish brain [33]. Future studies are required to investigate if *nmu* expression in the gut plays also a similar anorexigenic role in zebrafish and, since *crh* is not expressed in gut, how its potential regulatory function is mediated in the gut. In mice and rats, the anorexigenic effects of *nmu* are independent of leptin signal in the brain [130], but in the present study we found that *nmu* expression in the zebrafish gut is linked to leptin signal during refeeding. Finally, similar to our previous observations in the brain [33], the strongest divergence in expression of appetite-regulating genes between the genotypes in the gut were found during fasting.

## Conclusions

The study revealed that most brain-expressed orexigenic and anorexigenic genes are also expressed in the zebrafish gut. Unlike in the brain, changes in feeding conditions do not affect the expression of orexigenic genes in the zebrafish gut, as only *hcrt* showed differential expression in the gut of zebrafish with impaired leptin signal. On the contrary, among the anorexigenic genes, several genes were differentially expressed in the gut of wild-type and *lepr* mutant zebrafish and significant differences were found under different feeding conditions. Taken together, based on the results of the present study, we propose a potential role of the gastrointestinal system in the regulation of feeding in zebrafish. Future functional studies are required to unravel molecular cross-talks between brain and gut in regulation of feeding behaviour in zebrafish.

## Acknowledgements

The authors thank the Genome Engineering Zebrafish Facility at Uppsala University for its responsible management of our fish. Special thanks to Chrysoula Zalamitsou for her technical assistance through the study. The work was supported by the Carl Trygger Foundation for scientific research (CTS 16:413).

## Author Contributions

MS, MB and EPA designed the study. MB performed the fish breeding, feeding experiments, tissue sampling and RNA extraction. EPA designed the qPCR primers and studied literature for selection of candidate genes. EPA, JC and ET conducted the laboratory work for the gene expression step. EPA analysed the gene expression data and prepared the graphs. EPA, ET and MS wrote the manuscript. All authors reviewed the manuscript and approved its content.

## Ethical approval

The fish handling procedures were approved by the Swedish Ethical Committee on Animal Research in Uppsala (permit C10/16). All methods were carried out in accordance with the guidelines and regulations of the Swedish Ethical Committee on Animal Research in Uppsala.

## Data availability

All the data represented in this study are provided within the main manuscript or in the supplementary materials.

